# H7 influenza A viruses bind sialyl-LewisX, a potential intermediate receptor between species

**DOI:** 10.1101/2023.12.15.571923

**Authors:** Cindy M. Spruit, Diana I. Palme, Tiehai Li, María Ríos Carrasco, Alba Gabarroca García, Igor R. Sweet, Maryna Kuryshko, Joshua C. L. Maliepaard, Karli R. Reiding, David Scheibner, Geert-Jan Boons, Elsayed M. Abdelwhab, Robert P. de Vries

**Affiliations:** Department of Chemical Biology & Drug Discovery, Utrecht Institute for Pharmaceutical Sciences, Utrecht University, Utrecht, The Netherlands; Institute of Molecular Virology and Cell Biology, Friedrich-Loeffler-Institut, Federal Research Institute for Animal Health, Südufer 10, 17493, Greifswald, Insel Riems, Germany; Complex Carbohydrate Research Center, University of Georgia, Athens, GA, United States of America; Shanghai Institute of Materia Medica, Chinese Academy of Sciences, Shanghai, China; Biomolecular Mass Spectrometry and Proteomics, Bijvoet Center for Biomolecular Research and Utrecht Institute of Pharmaceutical Sciences, Utrecht University, Padualaan 8, 3584CH, Utrecht, The Netherlands

**Keywords:** Influenza A virus, hemagglutinin, interspecies transmission, sialyl-LewisX, NeuGc

## Abstract

Influenza A viruses (IAVs) can overcome species barriers by adaptation of the receptor binding site of the hemagglutinin (HA). To initiate infection, HAs bind to glycan receptors with terminal sialic acids, which are either *N*-acetylneuraminic acid (NeuAc) or *N*-glycolylneuraminic acid (NeuGc), the latter is mainly found in horses and pigs but not in birds and humans. We investigated the influence of previously identified equine NeuGc-adapting mutations (S128T, I130V, A135E, T189A, and K193R) in avian H7 IAVs *in vitro* and *in vivo.* We observed that these mutations negatively affected viral replication in chicken cells, but not in duck cells, and positively affected replication in horse cells. *In vivo*, the mutations reduced virus virulence and mortality in chickens. Ducks excreted high viral loads for a longer time than chickens, although they appeared clinically healthy. To elucidate why chickens and ducks were infected by these viruses despite the absence of NeuGc, we re-evaluated the receptor binding of H7 HAs using glycan microarray and flow cytometry studies. This revealed that mutated avian H7 HAs also bound to α2,3-linked NeuAc and sialyl-LewisX, which have an additional fucose moiety in their terminal epitope, explaining why infection of ducks and chickens was possible. Interestingly, the α2,3-linked NeuAc and sialyl-LewisX epitopes were only bound when presented on tri-antennary *N*-glycans, emphasizing the importance of investigating the fine receptor specificities of IAVs. In conclusion, the binding of NeuGc-adapted H7 IAV to sialyl-LewisX enables viral replication and shedding by chickens and ducks, potentially facilitating interspecies transmission of equine-adapted H7 IAVs. (249 words)

**Importance:** Influenza A viruses cause millions of deaths and illness in birds and mammals each year. The viral surface protein hemagglutinin initiates infection by binding to host cell terminal sialic acids. Hemagglutinin adaptations affect the binding affinity to these sialic acids and therefore the potential host species targeted. While avian and human IAVs tend to bind *N*-acetylneuraminic acid (a form of sialic acid), equine H7 viruses prefer binding to *N*-glycolylneuraminic acid (NeuGc). To better understand the function of NeuGc-specific adaptations in hemagglutinin and to elucidate interspecies transmission potential NeuGc-adapted viruses, we evaluated the effects of NeuGc-specific mutations in avian H7 viruses in chickens and ducks, important economic hosts and reservoir birds, respectively. We also examined the impact on viral replication and found a binding affinity to sialyl-LewisX, another terminal epitope. These findings are important as they contribute to the understanding of the role of sialyl-LewisX in avian influenza infection. (148 words)

## Introduction

Influenza A viruses (IAVs) are a member of the virus family *Orthomyxoviridae* and their proteins are encoded on eight single-stranded negative-sense RNA segments with a total length of 12-14kb (1). The enveloped virion of IAVs is coated with the surface proteins hemagglutinin (HA) and neuraminidase, which allow the classification into different subtypes (HxNx). IAVs infect a variety of avian and mammalian species, including humans, pigs, and horses (2). The natural reservoirs for IAVs are wild waterfowl, but transmission from ducks to other susceptible avian and mammalian species is frequent (3, 4). Avian IAV infection in wild birds is often asymptomatic, due to the coevolution of IAV and wild birds (5). However, in poultry low pathogenicity avian influenza viruses (LPAIV) can evolve into high pathogenicity avian influenza virus (HPAIV) causing mortality rates up to 100% in infected flocks. One of the key determinants for the virulence and pathogenicity of HPAIV is the acquisition of a multibasic cleavage site in the HA, which is most common in H5 and H7 IAVs (6).

High pathogenicity H7 IAVs are occasionally transmitted to humans and other mammalian species (7–9). Equine H7N7 influenza A viruses contain a multibasic cleavage site and are suspected to have originated from an avian H7 ancestor virus from an HPAIV outbreak in poultry (10). Furthermore, reassortant viruses with the equine H7N7 HA and other genes from a chicken H5N2 IAV were shown to be lethal in chickens (11). Nowadays, equine H7N7 viruses are thought to be extinct, leaving equine H3N8 as the only active circulating equine influenza virus (12, 13). The presence of H7 IAVs in different species emphasizes the relevance of further investigating these viruses and their interspecies transmission.

Overcoming host species barriers and establishing species-specific influenza strains involves the accumulation of point mutations during adaptation (14–17). The main host species barrier of IAVs is the receptor binding specificity of HA to terminal sialic acid (Sia) epitopes on the host cell surface, which is important for virus uptake into the cell (18). Receptor binding of HAs is strain-specific and has co-evolved with receptors found in the respiratory and/or intestinal tract of susceptible host species. Therefore, avian influenza viruses (AIV) bind preferentially to α2,3-linked Sia, whereas human-adapted strains prefer α2,6-linked Sia receptors (7, 19–21).

Besides the glycosidic linkage, the host cell receptor’s type of terminating Sia and the underlying glycan structures are important factors in IAV receptor binding properties and host range (14, 22–25). Unlike the majority of IAV, which predominantly bind to glycans with a terminal *N*-acetylneuraminic acid (NeuAc), equine H7N7 IAV predominantly bind to the *N*-glycolylneuraminic acid (NeuGc) (26, 27). Levels of NeuGc are variably present in the respiratory tract of most mammalian species, especially horses and pigs. However, no NeuGc is expressed in, among others, birds, humans, and ferrets (28–32). Previously, we identified five mutations S128T, I130V, A135E, T189A, and K193R in the receptor binding site (RBS), based on an equine H7N7 virus, that switched avian H7 IAVs from binding NeuAc to NeuGc (33).

In this study, we examined the impact of the equine NeuGc-adapting mutations (S128T, I130V, A135E, T189A, and K193R) on avian H7 IAVs both *in vitro* and *in vivo*, with particular emphasis on economically important poultry (chickens) and natural reservoir bird (ducks). The mutated viruses showed reduced replication in chicken cells, however, the replication in duck cells remained unaffected. On the other hand, viral replication in horse cells was increased. *In vivo*, the NeuGc-adapted viruses showed reduced mortality and virulence in infected chickens compared to the WT HPAIV, but the viral distribution between chicken organs was mostly unaffected by the mutations. However, virus shedding was higher in cloacal swab samples and ducks excreted high viral loads for a longer time than chickens, although they did not show symptoms of disease. Avian wild-type and mutant H7 hemagglutinins bound to both α2,3-linked NeuAc and sialyl-LewisX epitopes (an α2,3-linked NeuAc presented on an *N*-acetyllactosamine (LacNAc) with an additional fucose moiety α1,3-linked to the *N*-acetylglucosamine of the LacNAc), but only when presented in complex *N*-glycans. These findings improve the understanding of equine-specific adaptations in avian H7 receptor binding, viral replication, and pathogenicity while assessing the potential of interspecies transmission of these viruses.

## Results

### NeuGc-specific mutations have differential effects in chicken, duck, and horse cells

Previously, we investigated the molecular determinants for binding of avian H7 IAVs to NeuGc and found that five amino acids that are abundant in equine H7 viruses (128T, 130V, 135E, 189A, and 193R) were responsible for binding to NeuGc, a common sialic acid in horses (33). Curiously, we observed that these mutations in the H7 HA of A/turkey/Italy/214845/02 switched the receptor binding specificity from α2,3-linked NeuAc to α2,3-linked NeuGc, but did not cause a loss of binding to chicken trachea and erythrocytes, which do not contain NeuGc (29, 31). This observation raised the question of whether the infection capabilities of avian viruses with these equine NeuGc-specific mutations would be affected.

To investigate whether the NeuGc-specific mutations would affect the viral fitness of an avian virus *in vitro*, we rescued A/chicken/Germany/R28/2003 as a wild-type (WT) H7N7 HPAIV (designated H7N7_avHA) and a mutant of this virus carrying the five NeuGc-specific mutations in the HA (designated H7N7_5eqHA). The RBS of H7N7_avHA is identical to the RBS of A/turkey/Italy/214845/02, which we used in our previous publication (33). Sequence analysis of avian and equine H7 sequences showed that all five amino acid residues are highly conserved in equine H7 IAVs (97-100%), but also naturally occur in some of the analyzed avian H7 HA sequences (Table S1).

The impact of the equine-specific residues on cell-to-cell spread and viral growth kinetics was investigated in various cell lines (Fig. 1A-E). The viruses’ ability to spread from one cell to another was analyzed in a plaque assay using MDCKII cells, which are the most commonly used cells for IAV replication assays (34). The NeuGc-specific residues significantly reduced the intercellular spread at 72 hours post-infection (hpi) in MDCKII cells (Fig. 1A) and the replication in these cells (Fig. 1B). The NeuGc-specific mutations significantly increased viral replication in equine lung cells (PLU-R) and equine epidermal cells (E.Derm) 24 hpi (Fig. 1C). In contrast, H7N7_5eqHA replication in chicken fibroblasts, primary CEK cells, and SPF eggs was significantly reduced compared to WT H7N7_avHA (Fig. 1D). However, no significant differences between replication of H7N7_avHA and H7N7_5eqHA were observed in duck embryo fibroblast cells 24 hpi (Fig. 1E). These findings indicate that the equine-specific amino acid residues in the RBS of H7 HPAIV reduced cell-to-cell spread and viral replication in a host-dependent manner. They increased replication of the avian virus in horse cells and reduced replication in chicken cells, but replication in duck cells was unaffected.

**Figure 1.**
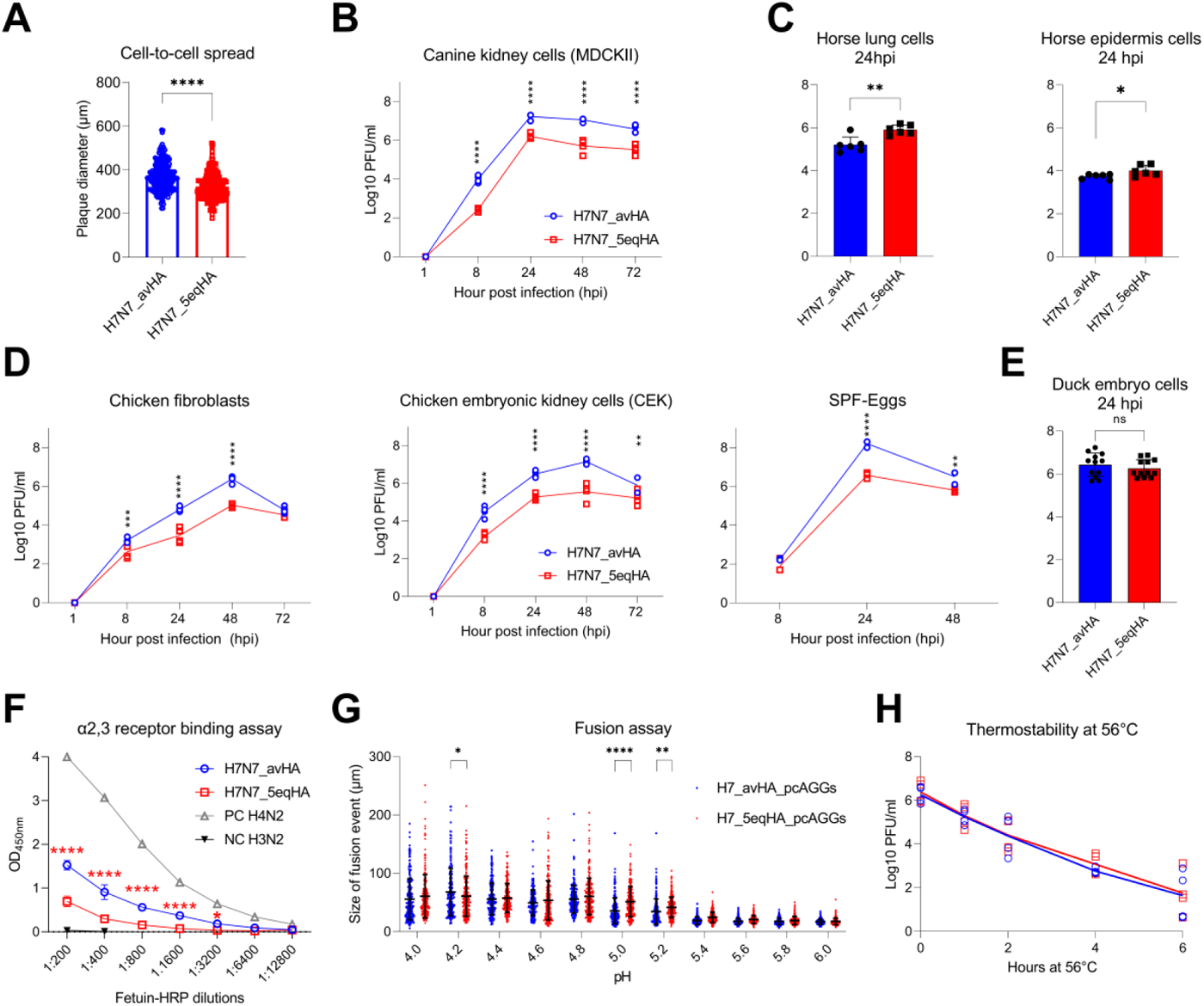
*In vitro* characterization of WT (H7N7_avHA) and mutant (H7N7_5eqHA) A/chicken/Germany/R28/2003 H7N7 viruses. (A) Cell-to-cell spread was investigated by measuring the diameter of about 100 plaques in MDCKII cells. (B) Viral replication at indicated time points was assessed in MDCKII, (C) horse lung and horse epidermal cells, (D) chicken fibroblasts (DF-1), primary chicken cells (CEK), SPF embryonated chicken eggs (ECE), and (E) in duck embryo fibroblast cells. (F) The receptor binding affinity to α2,3-linked NeuAc was measured using α2,3-Sia fetuin substrate. Human H3N2 virus (specific for α2,6-linked NeuAc) was used as a negative control (NC). An avian H4N2 virus (specific for α2,3-linked NeuAc) was used as a positive control (PC). Shown are representative results calculated as means and standard deviations of three independent experiments, each was run in duplicates. (G) pH-dependent activation of HA in a fusion assay was measured after transfection of quail cells (QM-9) with pCAGGS protein expression vector containing the HA of H7N7_avHA or H7N7_5eqHA. Cells were simultaneously transfected with pCAGGS carrying eGFP to facilitate the evaluation of the assay. Cell fusion was triggered 24 hpi with PBS of different pH values for two minutes. The diameter of syncytia was measured using Eclipse Ti-S with software NIS-Elements, version 4.0; Nikon. (H) The thermostability of viruses was measured in duplicates and repeated twice. The reduction in virus infectivity at indicated time points was assessed by titration of heated viruses using a plaque test in MDCKII cells and expressed as plaque-forming units per ml (Log10 PFU/ml). All results are expressed as means and standard deviations of at least two independent experiments run in duplicates. Asterisks indicate statistical significance based on p values: * ≤ 0.05, ** ≤ 0.01, *** ≤ 0.001, **** ≤ 0.0001.

### NeuGc-specific mutations reduced the binding affinity to α2,3-NeuAc without a significant impact on the pH-dependent HA activation and thermostability

*In vitro* characterization of generated H7N7 viruses in cell culture revealed that NeuGc-specific mutations affected viral replication and spread in a host-dependent manner (Fig. 1A-E). To ascertain whether these mutations have an influence on the viruses’ biological properties and thus on viral replication, the receptor binding properties, thermostability, and pH activation were tested.

To assess whether the introduction of the NeuGc-specific mutations in the RBS changed the binding affinity to α2,3-NeuAc, a solid-phase assay using α2,3-linked fetuin as a substrate was performed (Fig.1F) as previously described (26, 35). An avian H4N2 virus was used as a positive control for α2,3-NeuAc binding and a human H3N2 IAV was used as a negative control. We observed that H7N7_5eqHA had a significantly lower binding affinity to α2,3-NeuAc than the WT H7N7_avHA virus (Fig. 1F), which is in accordance with our previously obtained results (33). In addition to affecting receptor binding properties, mutations in HA1 may affect pH activation of hemagglutinin and subsequently affect the replication of viruses such as AIV H5N1 [53], although there is limited knowledge of how it affects AIV H7N7. Therefore, we assessed the potential influence of the five mutations on the pH-dependent fusion-HA activation by measuring the diameter of cell-to-cell fusion after transfection of avian cells (QM9) with protein expression vectors (pCAGGS) carrying HA from H7N7_avHA (H7_avHA_pcAGGs) or H7N7_5eqHA (H7_5eqHA_pcAGGs) (Fig. 1G). Both hemagglutinins were activated at a broad range of pH values from 4.0 to 6.0. However, the fusion efficiency of H7_5eqHA_pcAGGs in QM9 cells at a pH of 5.0 and 5.2 was significantly higher than that of H7_avHA_pcAGGs. Interestingly, the pH-dependent activation of H7_avHA_pcAGGs was found to be significantly higher at a pH value of 4.2 than H7_5eqHA_pcAGGs. The size of fusion events at other pH values was comparable (Fig. 1G).

The thermostability of the HA is known to be linked to virulence in different influenza strains (36). Therefore, we evaluated the thermostability of the two viruses at 56°C for 2, 4, and 6 hours, a standard treatment for enveloped viruses (37). Both viruses lost infectivity at comparable levels indicating that the introduction of equine-specific amino acids did not affect the thermostability of the HPAIV (Fig. 1H). In conclusion, the NeuGc-specific mutations reduced the replication of this H7N7 HPAIV in the chicken cells probably due to reduced binding affinity to the 2,3-NeuAc without significantly impacting the HA pH-dependent activation and thermostability of the viruses.

### The NeuGc-specific residues significantly reduced virulence in infected chickens, but had no impact on virus virulence in ducks

Since the NeuGc-specific mutations had an impact on receptor binding, cell-to-cell transmission, and viral replication in chicken, but not duck cells (Fig. 1), these mutations potentially also affect the viruses *in vivo.* Therefore, we performed an animal experiment in chickens, from which the H7N7_avHA was originally isolated and which are economically crucial hosts, and ducks as the natural reservoir species of AIVs.

Nine SPF chickens and ten Pekin ducks were infected by intravenous (IV) or intramuscular (IM) injection, respectively with H7N7_avHA or H7N7_5eqHA to assess the viral pathogenicity index (PI) according to the WOAH standard (38). All ducks infected IM with H7N7_avHA or H7N7_5eqHA survived the animal experiment and showed no clinical disorders (Table 1). Nevertheless, all ducks seroconverted indicating a successful infection. The intramuscular pathogenicity index (IMPI) for H7N7_avHA and H7N7_5eqHA in ducks was determined to be 0.0. Conversely, chickens that were IV-infected with H7N7_avHA died within a mean time of death (MDT) of 4.6 days post-infection (dpi). All chickens displayed clinical signs of infection and an intravenous pathogenicity index (IVPI) of 2.4 was calculated. H7N7_avHA is therefore classified as an HPAIV according to the WOAH classification (IVPI > 1.2 indicates an AIV as HPAIV). Interestingly, the introduction of the five equine mutations into the avian HA reduced mortality to 3 out of 9 IV-infected chickens with H7N7_5eqHA, but all chickens exhibited transient mild to moderate clinical signs. Notably, IV-infected chickens with H7N7_5eqHA died earlier compared to those IV-infected with H7N7_avHA, with an MDT of 3.0 days (Table 1), although the differences in MDT of both groups were not statistically significant. Nevertheless, the IVPI of H7N7_5eqHA inoculated chickens was determined to be 1.5, which is still classified as HPAIV. All remaining chickens subsequently developed antibodies.

**Table 1.** The pathogenicity indices of H7N7 viruses after the injection of chickens and ducks. The table shows the mortality, morbidity, and seroconversion of intravenous infected (IV) chickens and intramuscular infected (IM) Pekin ducks. In addition, the calculated mean time of death (MDT) in days post-infection (dpi) and intravenous (IVPI) or intramuscular (IMPI) pathogenicity indices are shown.

| Virus | Animal | Group | Mortality | Mean time of death (MDT) | Morbidity | Pathogenicity index (PI) | Seropositive birds/total birds |
| --- | --- | --- | --- | --- | --- | --- | --- |
| **H7N7_avHA** | Chickens | IV infected | 9/9 | 4.6 | 9/9 | IVPI: 2.4 | n.a |
|  | Ducks | IM infected | 0/10 | n.a. * | 0/10 | IMPI: 0.0 | 10/10 |
| **H7N7_5eqHA** | Chickens | IV infected | 3/10 | 3.0 | 10/10 | IVPI: 1.5 | 7/7 |
|  | Ducks | IM infected | 0/10 | n.a. * | 0/10 | IMPI: 0.0 | 10/10 |

\*n.a. = not applicable

In the second animal experiment, we wanted to mimic the natural course of infection. Therefore, ten chickens and ten ducks were inoculated by the oculonasal (ON) route. Furthermore, at 1 dpi five chickens and five ducks were added to each group to assess chicken-to-chicken or duck-to-duck transmission. All chickens primarily inoculated with H7N7_avHA died within an MDT of 5.7 dpi, with an average CS of 1.7 (Fig. 2A, Table 2). Only one contact chicken died on the eighth dpi in the avian H7N7 (H7N7_avHA) infected group. However, all contact chickens displayed signs of morbidity (Table 2). Six out of ten chickens infected with H7N7_5eqHA died with an MDT of 4.5 days. In this group, three out of five contact chickens died with MDT of 7.3 days, and the two remaining chickens showed transient mild to moderate clinical signs. The average PI was 1.4 for the primarily inoculated chickens (Fig. 2A, Table 2). Conversely, and similar to the IM-injected ducks, neither the ON-inoculated nor the contact ducks in either group displayed any signs of illness. All ducks survived until the end of the animal trial (Table 2, Fig. 2B). Seropositive results were recorded for all remaining chickens and ducks using an anti-NP ELISA (Table 2). These findings confirm the results from the IVPI and IMPI-infected birds, as H7N7_5eqHA is less lethal in chickens than H7N7_avHA and ducks are not affected at all. Taken together, and regardless of the infection routes, the equine-adaptive mutations reduced HPAIV H7N7 virulence in chickens, while ducks were clinically resistant to both viruses.

**Figure 2.**
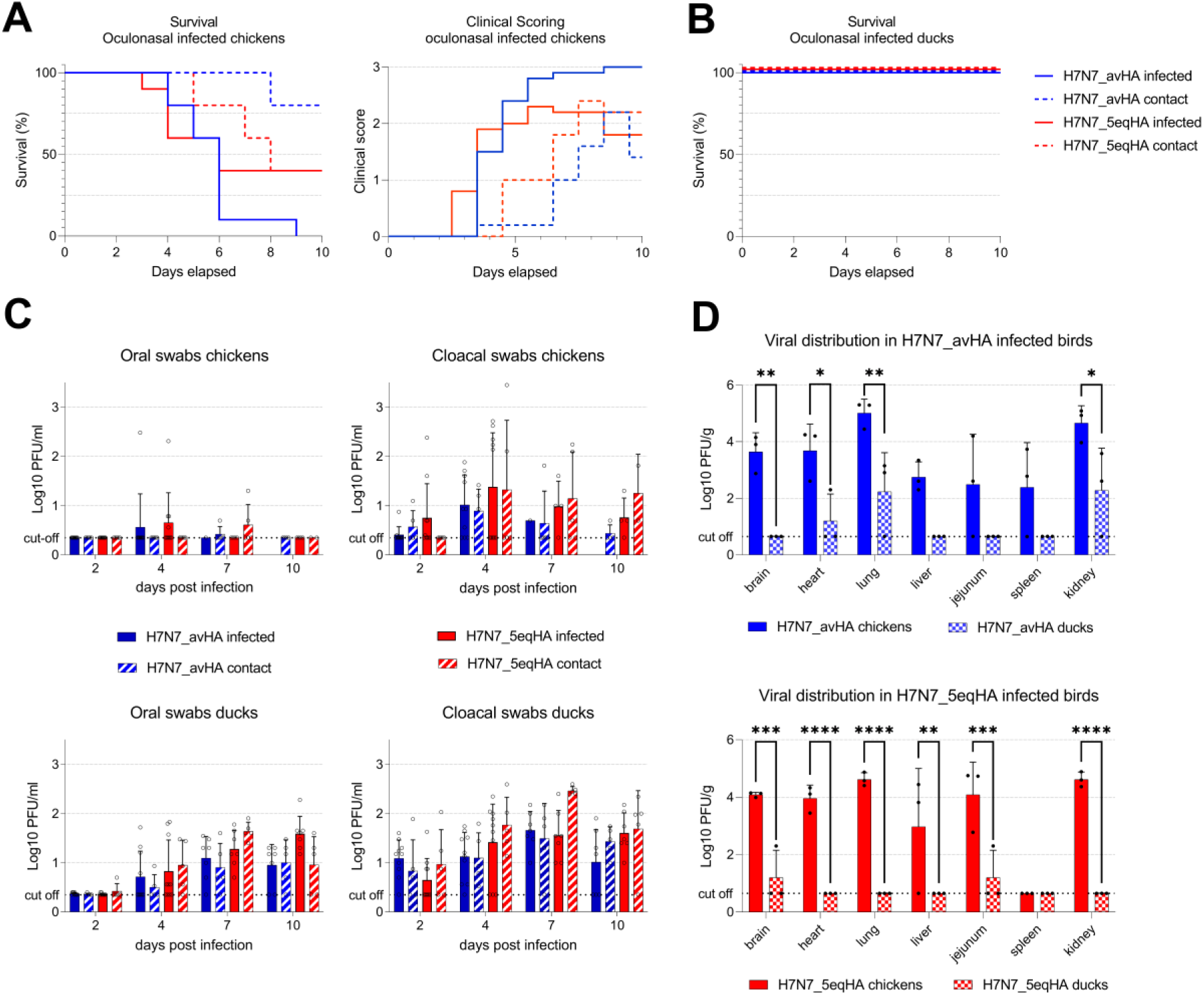
*In vivo* characterization of WT (H7N7_avHA) and mutant (H7N7_5eqHA) A/chicken/Germany/R28/2003 H7N7 virus. (A) Survival and clinical score of ON infected chickens throughout the animal experiment. (B) Survival curve of ON infected Pekin ducks. (C) Analysis of oral and cloacal swab samples taken from chickens and ducks in plaque tests expressed as Log10 PFU/ml. (D) The viral distribution in duck and chicken organs was analyzed in plaque tests and expressed as PFU/gram. Asterisks indicate statistical significance based on p values * ≤ 0.05, ** ≤ 0.01, *** ≤ 0.001, **** ≤ 0.0001. ns = not significant. Dashed lines indicate the predicted detection limit of the plaque assay (cut-off).

**Table 2.** *In vivo* data from oculonasal infected chickens and Pekin ducks. The table shows the sick (morbidity) and dead (mortality) animals per group, as well as the results from sera analysis in a competitive NP ELISA. Calculated mean time to death (MDT) in days post-infection as well as the clinical score (CS) are shown.

| Virus | Animal | Group | Mortality | Mean time of death (MDT) | Morbidity | Clinical Score (CS) | Seropositive birds /total birds |
| --- | --- | --- | --- | --- | --- | --- | --- |
| **H7N7_**<br>**avHA** | Chickens | ON infected | 10/10 | 5.7 | 10/10 | 1.7 | n.a |
|  |  | ON Contact | 1/5 | 8.0 | 5/5 | 0.6 | 4/5 |
|  | Ducks | ON infected | 0/10 | n.a.* | 0/10 | 0.0 | 7/7 |
|  |  | ON Contact | 0/5 | n.a.* | 0/5 | 0.0 | 5/5 |
| **H7N7_**<br>**5eqHA** | Chickens | ON infected | 6/10 | 4.5 | 10/10 | 1.4 | 4/4 |
|  |  | ON Contact | 3/5 | 7.3 | 5/5 | 1.0 | 2/2 |
|  | Ducks | ON infected | 0/10 | n.a.* | 0/10 | 0.0 | 7/7 |
|  |  | ON Contact | 0/5 | n.a.* | 0/5 | 0.0 | 5/5 |

\*n.a. = not applicable

### The NeuGc-specific residues did not affect virus replication or excretion in chickens, but ducks are potentially silent spreaders of H7N7 viruses

We further determined the effect of the five equine mutations on viral loads in swab and organ samples obtained from ON-inoculated birds and their contacts. Oral and cloacal swabs collected 2, 4, 7, and 10 dpi were tested using plaque assay. In chickens and ducks, the level of virus shedding from the cloacal route was higher compared to the oral route, although the differences were not statistically significant (Fig. 2C). Interestingly, ducks excreted high viral loads by the oral and fecal routes for a longer time than chickens. Moreover, the viral distribution in different organs (brain, heart, lungs, liver, jejunum, spleen, and kidneys) obtained 4 dpi from three birds of each group was broader in chickens than in ducks. In H7N7_avHA-infected chickens, the viral load was significantly higher in the brain, heart, liver, and kidneys compared to infected ducks (Fig. 2D). Similarly, in H7N7_5eqHA-infected chickens, the viral load was significantly higher in all organs except the spleen compared to ducks.

In conclusion, we observed increased levels of viral shedding via the cloacal routes and prolonged shedding of viruses in ducks compared to chickens. The viral distribution in chicken organs was broader than in ducks, which may explain the high mortality in chickens. These findings raise concerns about the potential spread of H7N7 viruses, particularly by ducks as silent spreaders. The high and prolonged shedding of the H7 viruses in ducks, even without exhibiting clinical symptoms, pose a potential risk for their reintroduction to hosts that have a high presence of NeuGc, like pigs and horses.

### Avian H7 IAVs bind both α2,3-linked NeuAc and sialyl-LewisX epitopes

The animal experiments showed that both chickens and ducks were infected by the NeuGc-specific H7 viruses (Fig. 2), although both species are known to not express NeuGc (29, 31). This strongly suggests that in our previous research, in which we observed NeuGc-specificity of this mutant (S128T, I130V, A135E, T189A, and K193R) H7 HA (33), we overlooked the binding to one of the many other glycans that may be present in nature. We hypothesized that sialyl-LewisX (sLe^x^) epitopes are important as they have recently been shown to be involved in H7 IAV infection (39, 40) and are bound by nearly all IAV subtypes (41–49). The sLe^X^ epitope consists of an α2,3-linked NeuAc presented on an *N*-acetyllactosamine (LacNAc) with an additional fucose moiety α1,3-linked to the *N*-acetylglucosamine (GlcNAc) of the LacNAc. The sLe^X^ epitopes are present in some species and tissues, such as chicken trachea and colon, guinea fowl trachea, turkey respiratory tract, and human lung (45, 50–55). Most research investigating binding to the sLe^X^ epitope has been performed using a tetrasaccharide sLe^X^ epitope, due to a lack of biologically relevant glycans.

Here, we investigated which exact glycans are bound by WT and mutant avian H7 HAs to explain how chickens and ducks are infected by NeuGc-specific H7 viruses. Since the complex glycan structure can influence receptor binding (14, 22–25), we here focused on biologically relevant complex *N*-glycans presenting α2,3-linked NeuAc, α2,3-linked NeuGc, α2,6-linked NeuAc, and sLe^X^ epitopes (Fig. S1). These glycans were printed on glass slides and after incubation of the slides with the HAs and fluorescent secondary antibodies, the glycan-HA binding was evaluated.

Whereas the WT HA of A/turkey/Italy/214845/02 (H7tu) previously only showed binding to glycans terminating in α2,3-linked NeuAc (33), here we observed a strong preference for tri-antennary *N*-glycans presenting at least one sLe^X^ epitope (glycans **25**-**28**, Fig. 3). Both glycans solely presenting sLe^X^ epitopes (**27**, **28**) and glycans presenting a sLe^X^ on one arm and α2,3-linked NeuAc on the other two arms (**25**, **26**) were bound. For the latter, it cannot be distinguished whether binding is caused by the sLe^X^ or the α2,3-linked NeuAc. Interestingly, binding to sLe^X^ epitopes presented on linear glycans (**7–9**) was not observed. Furthermore, steric hindrance due to the presence of α2,6-linked NeuAc on two arms, besides the sLe^X^ on one arm, appeared to be present, as glycan **23** and **24** were not bound. We also showed that the mutant H7tu HA bound both glycans terminating in α2,3-linked NeuGc, as well as sLe^X^-presenting tri-antennary *N*-glycans (Fig. 3). In conclusion, the observed binding to sLe^X^ (Fig. 3) showed that the previously studied mutant avian H7 HAs were not strictly specific for NeuGc and may explain why these HAs bound to chicken tissue and erythrocytes previously (33). This sLe^X^-binding possibly also explains why ducks and chickens could be infected by the NeuGc-adapted avian H7 virus (Fig. 2).

**Figure 3.**
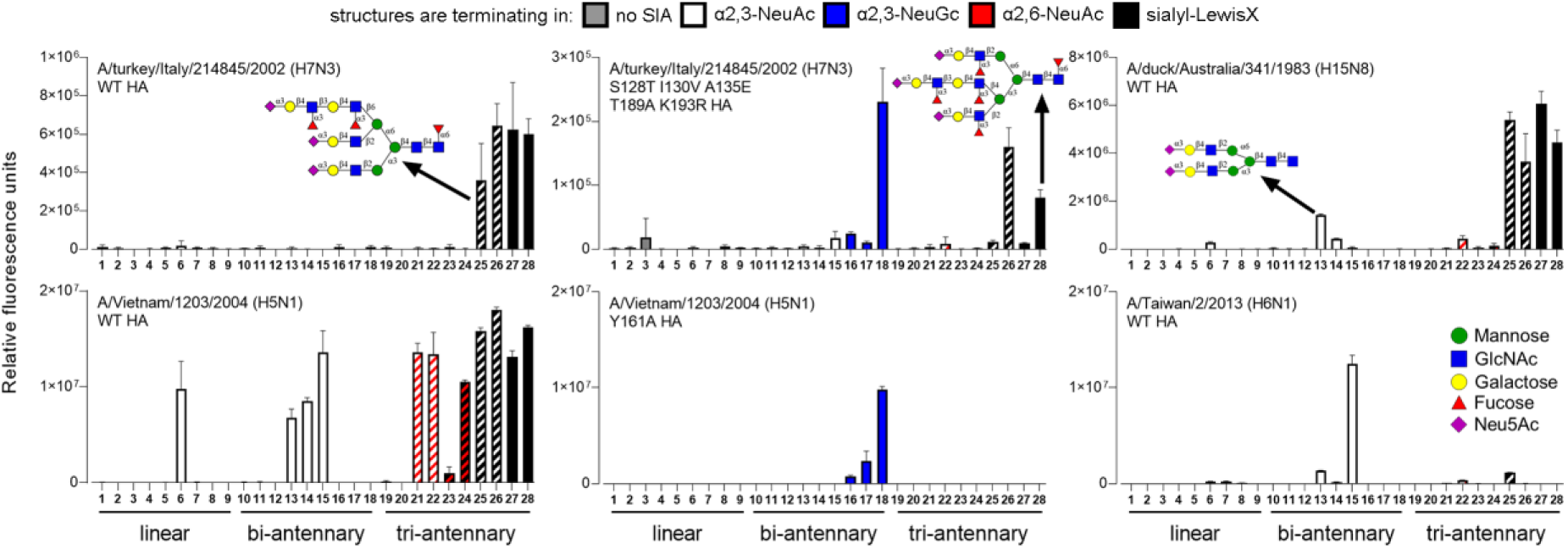
Avian H7 HAs bind to sialyl-LewisX epitopes. Synthetic glycans were used to assess the receptor binding of the IAV HAs (A/turkey/Italy/214845/2002 H7, A/duck/Australia/341/1983 H15, A/Vietnam/1203/2004 H5, and A/Taiwan/2/2013 H6). The glycans were terminating in galactose (no SIA, grey), α2,3-linked NeuAc (white), α2,3-linked NeuGc (blue), α2,6-linked NeuAc (red), or sialyl-LewisX (black). Bars with two colors indicate glycans terminating in different epitopes on different arms. Fig. S1 presents all structures that are present on the array. Bars represent the mean ± SD (n=4).

Furthermore, we examined the receptor specificity of the H15 HA of A/duck/Australia/341/1983, the closest related subtype to H7. The H15 HA showed a similar binding phenotype to the H7tu HA and bound α2,3-linked NeuAc and sLe^X^, while structures presenting α2,6-linked NeuAc on the other arms were not bound (Fig. 3). Interestingly, H15 HAs were previously not known to bind to sLe^X^ epitopes. The steric hindrance due to α2,6-linked NeuAc appeared to be specific for the H7 and H15 HAs, since the WT H5 HA from A/Vietnam/1203/2004 (H5VN) showed binding to all tri-antennary *N*-glycans presenting at least one sLe^X^ epitope, regardless of the terminal epitopes presented on the other arms (Fig. 3). Consistent with the results from H7 and H15 HAs, the WT H5VN HA only bound to sLe^X^ epitopes when presented on tri-antennary *N*-glycans, and not linear glycans, possibly due to a multivalency effect because of the high density of binding epitopes in one glycan. Furthermore, an HA that we previously used as a control for specific binding of NeuGc (the Y161A HA mutant A/Vietnam/1203/2004 H5N1) bound specifically to NeuGc and not sLe^X^ (Fig. 3). Interestingly, the H6 HA from A/Taiwan/2/2013 was strictly specific for α2,3-linked NeuAc and did not bind to glycans that present other epitopes on the other arms (**25** and **26**). The results showed that the fine receptor specificity is highly dependent on the IAV and the exact complex glycan structure.

Nevertheless, the binding of the H7 HAs to sLe^X^ epitopes does not explain the infection in ducks, since ducks are generally assumed not to present NeuGc nor sLe^X^ epitopes (29, 31, 50). A tissue stain using anti-sialyl-LewisX antibodies revealed that, indeed, no sLe^X^ epitopes were found on duck colon and tracheal tissues (Fig. 4A). Therefore, we aimed to investigate whether the binding of these avian H7 HAs was truly dependent on sLe^X^ epitopes. We performed assays using a fucosidase E1_10125 from *Ruminococcus gnavus* E1, which specifically cleaves the fucose moiety from the sLe^X^ epitope (56) (Fig. 4B). The anti-sialyl-LewisX antibody showed that no more sLe^X^ epitopes remained on the chicken trachea after fucosidase treatment. However, the binding of both WT and mutant H7tu (Fig. 4B) HA to chicken trachea remains unchanged after fucosidase treatment, indicating that α2,3-linked NeuAc is bound in a tissue section.

To investigate which exact glycans may have been involved in this binding to the chicken trachea, we treated the glycan microarray with fucosidase (Fig. 4C). Using an anti-sLe^X^ antibody that bound all sLe^X^-containing structures on the glycan microarray as a control, we indeed showed that all sLe^X^ epitopes were removed after fucosidase treatment of the array. This suggests that no sLe^X^ epitopes remained on the chicken trachea either after fucosidase treatment (Fig. 4B). As additional controls, we used the WT and R222K R227S mutant H5 HAs of A/chicken/Ibaraki/1/2005 (H5IBR), which are, respectively, specifically binding to sLe^X^ structures and α2,3-linked NeuAc (43). After fucosidase treatment, these WT and mutant H5IBR HA controls showed, respectively, no binding to sLe^X^ glycans and increased binding to sLe^X^ glycans (that are now converted to α2,3-linked NeuAc) (Fig. 4C). Interestingly, the H7tu WT HA still bound to structures **25** to **28** (of which the sLe^X^ is converted to α2,3-linked NeuAc) after fucosidase treatment, but not bi-antennary *N*-glycans presenting α2,3-linked NeuAc (**13**-**15**, Fig. S1, Fig. 4C), suggesting that tri-antennary *N*-glycans are preferred as receptors for H7tu. In conclusion, both α2,3-linked NeuAc and sLe^X^ epitopes presented on the tri-antennary *N*-glycans are bound efficiently by the H7tu WT HA.

To further investigate binding to sLe^X^ glycans, we used MDCK WT and MDCK-FUT cells. The latter overexpress the chicken fucosyltransferase genes *FUT3*, *FUT5*, and *FUT6* (50), which was expected to increase the amount of sLe^X^ epitopes that are presented on the cells. To investigate whether indeed increased amounts of sLe^X^ epitopes were presented on the cells, we employed mass spectrometry (MS) methods. We first investigated the released *N*-glycans from MDCK WT and FUT cells by HILIC-IMS-QTOF positive mode MS and found that MDCK-FUT cells presented a higher number of fucoses on sialylated *N*-glycans than MDCK WT cells (Fig. 4D, Table S2). To further investigate whether the fucoses were present in sLe^X^ epitopes, we analyzed the *N*-glycans using fragmentation in LC-MS/MS, which indeed showed a higher relative abundance of sLe^X^ fragments (oxonium ions of *m/z* 803.2928) on the MDCK-FUT cells (Fig. 4E). We then continued to use these MDCK WT and FUT cells in flow cytometry analysis. The controls (anti-sLe^X^ and the HA of A/Taiwan/2/2013 H6N1, which is specific for α2,3-linked NeuAc (Fig. 3)) showed that the amount of sLe^X^ on MDCK WT cells and the amount of α2,3-linked NeuAc on MDCK-FUT cells was very low (Fig. 4F). Surprisingly, the H5IBR HAs (WT and mutant) that were assumed to be specific for sLe^X^ and α2,3-linked NeuAc, respectively, bound well to both cell types. Similar to the result in the glycan microarray and tissue stains, the H7 (and H15) WT and mutant HAs bound to both cell types. In conclusion, both α2,3-linked NeuAc and sLe^X^ epitopes appear to be bound efficiently by the H7tu, but binding depends on the exact glycan structure.

**Figure 4.**
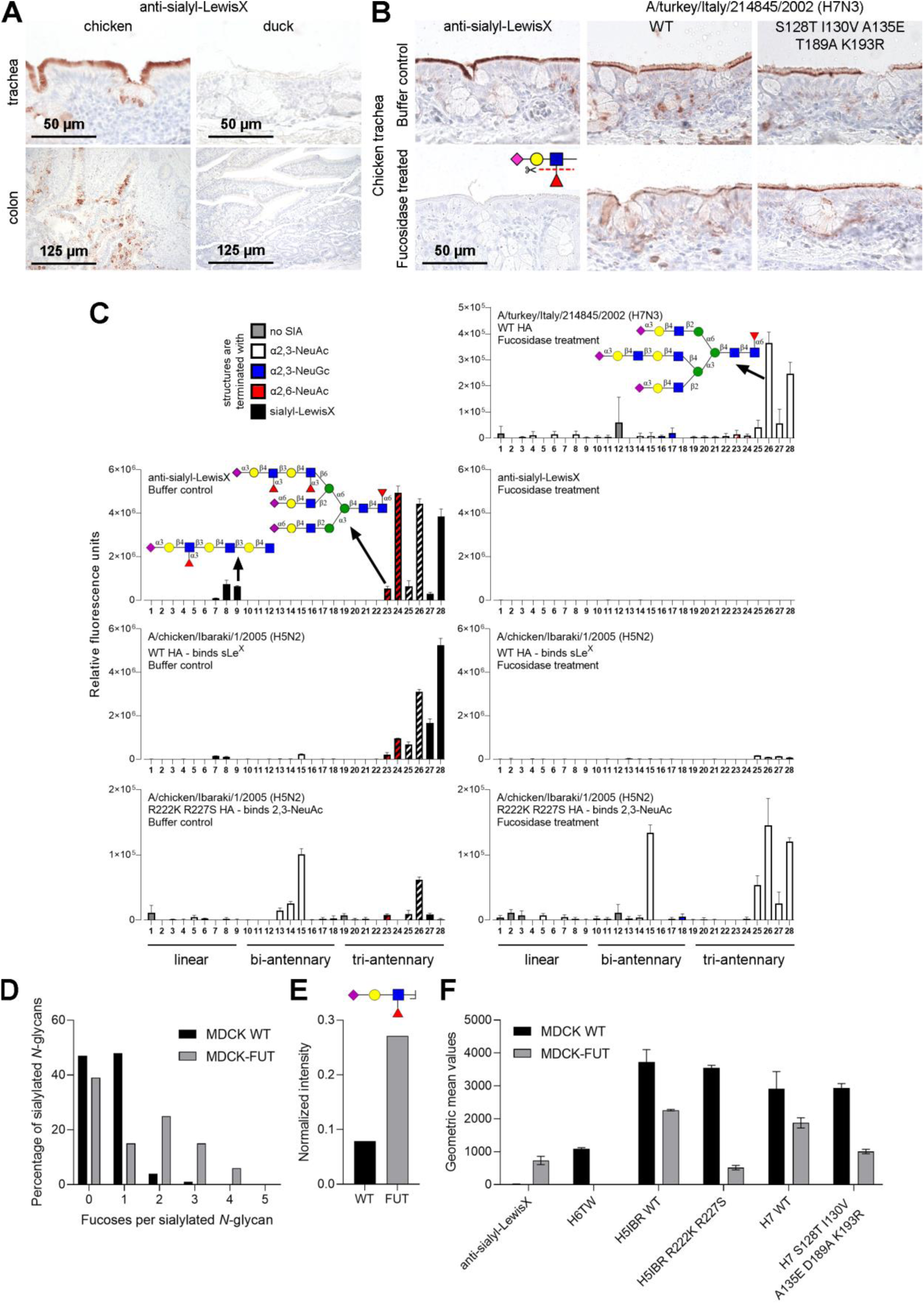
Avian H7 HAs bind both α2,3-linked NeuAc and sialyl-LewisX epitopes. (A) The presence of sialyl-LewisX epitopes on chicken and duck trachea and colon was investigated using anti-sialyl-LewisX antibodies. (B) The binding of anti-sialyl-LewisX antibodies and H7 HAs to chicken tracheal tissue (with and without fucosidase treatment) was assessed. Tissue binding was visualized using AEC staining. (C) Synthetic glycans (with and without fucosidase treatment) were used to assess the binding of the anti-sialyl-LewisX antibody, the WT H7 HA of A/turkey/Italy/214845/02, and H5 HAs of A/chicken/Ibaraki/1/2005. (D) The *N*-glycans of MDCK WT and MDCK-FUT cells were investigated using HILIC-IMS-QTOF positive mode mass spectrometry. The number of fucoses per sialylated *N*-glycan was analyzed for both cell types. Further analysis is presented in Table S2. (E) The *N*-glycans were further analyzed using LC-MS/MS. The oxonium ions of m/z 803.2928 (sLe^X^) were identified and normalized to the core fragments. Mean and standard errors (n=3) are shown. (F) Flow cytometry measurements were performed to assess the binding anti-sialyl-LewisX antibodies and HAs to MDCK WT and MDCK-FUT cells. Triplicate measurements were performed, of which the mean and standard deviation are displayed.

## Discussion

Here, we studied the effects of equine NeuGc-adapting mutations (S128T, I130V, A135E, T189A, and K193R) in avian H7 IAVs *in vitro* and *in vivo*. These viruses are potentially candidates for interspecies transmission between avian and mammalian species expressing NeuGc receptors. The insertion of equine NeuGc-adapting mutations resulted in stable and viable viruses and increased viral replication in horse cells. While the mutations reduced viral replication in chicken and dog cells, interestingly the replication in duck cells was not affected. *In vivo*, the NeuGc-adapting mutations not only reduced the pathogenicity index in intravenously infected chickens but also mortality and morbidity in oculonasal-infected chickens. In ducks, on the other hand, neither virus caused signs of illness or increased mortality. Nevertheless, ducks shed high amounts of virus for a longer time compared to chickens. Here, NeuGc-adapting mutations were not disadvantageous in viral shedding compared to the WT HPAIV. The NeuGc-adapted H7 was additionally found to bind α2,3-linked NeuAc and sialyl-LewisX (sLe^X^) epitopes, but only when these epitopes were presented on tri-antennary *N*-glycans. Binding to these epitopes explains why ducks and chickens could be infected and emphasizes the risk of interspecies transmission of H7 IAVs.

Although sLe^X^ epitopes were identified as potential receptors for the studied H7 viruses, it is currently unclear whether sLe^X^ is used in IAV infections as a functional receptor, or whether it has other functions. It has been suggested that the presence of sLe^X^ facilitates H7 IAV infection . If sLe^X^ binding is important in IAV infection, this may cause a species barrier or act as an intermediate receptor since some species and tissues, such as the chicken trachea and colon, guinea fowl trachea, turkey respiratory tract, and human lung present sLe^X^ epitopes (45, 50–55).

The molecular determinants for the binding of H7 viruses to sLe^X^ epitopes are currently unknown. Since we showed that not all IAVs bind to sLe^X^ epitopes, such as the H6 HA of A/Taiwan/2/2013 (Fig. 3), likely certain amino acids are responsible for the binding to sLe^X^. For H5 viruses, it was shown that mutations K222R/Q and S227R in the HA convert from binding to α2,3-linked NeuAc to sLe^X^ (43, 49, 51). Especially, the lysine at position 222 was shown to sterically hinder binding to sLe^X^ epitopes (57), while a glutamine or arginine at that position enables, potentially through a hydrogen bond, sLe^X^ binding. Indeed, the H7 viruses that were investigated here contain a glutamine (Q) at position 222, which is highly conserved in H7 viruses (51) and partially explains the binding to sLe^X^. However, position 227 is also a glutamine in the investigated H7 viruses, of which the effect on sLe^X^ binding is currently unknown. Elsewhere, the presence of a lysine at position 193 is reported to be important for sLe^X^ binding and, indeed, the H7tu contains a 193K (40). Additionally, amino acids at other positions, which have not been investigated yet, may also affect the binding to sLe^X^ epitopes.

Previously, IAVs from all subtypes, except H15, were shown to bind sLe^X^ epitopes (41–49). Using tri-antennary *N*-glycans presenting sLe^X^ epitopes, we here showed that also H15 IAVs, the closest related subtype to H7, are capable of binding sLe^X^ epitopes. Furthermore, the investigated avian H7 HAs were previously not known to bind to sLe^X^ epitopes, as binding was only observed when the epitopes were presented on tri-antennary *N*-glycans and not linear glycans. Additionally, the WT H7 HA appeared to bind stronger to α2,3-linked NeuAc when presented on tri-antennary *N*-glycans (sLe^X^ after fucosidase treatment) than bi-antennary *N*-glycans or linear glycans. The H7tu also appeared to bind stronger to tri-antennary *N*-glycans presenting the α2,3-linked NeuAc on an elongated MGAT4 arm (**26** and **28**) instead of an elongated MGAT5 arm (**25** and **27**) (Fig. 4C, Fig. S1), although this was not consistent throughout the replicates. Using these sLe^X^-presenting *N*-glycans in combination with other HAs may reveal the binding of more IAVs to sLe^X^ and the role of these epitopes in IAV infection. These observations highlight the relevance of looking beyond the terminal epitope and considering the fine receptor specificity when investigating IAV receptor binding.

The distribution and types of Sias are species-specific and variable throughout the respiratory tract of IAV-susceptible species (17). HA specificity is often adapted to the particular Sia receptors present in the host (14). Thus, interspecies transmission and establishment in a new host requires a successful adaptation of HA binding specificity to the new host environment as seen in equine H7N7 viruses originating from avian H7 viruses (10). Although equine H7N7 IAVs are thought to be extinct (12, 13), the amino acid residues coding for the NeuGc binding specificity persist in avian H7 sequences (Table S1), enabling a potential re-emergence of NeuGc-binding viruses. The avian H7 IAVs with equine-adapted mutations that we investigated not only bound to equine-specific NeuGc-receptors but were also able to replicate and infect avian hosts (Fig. 2). The viruses with equine-adapted NeuGc-specific mutations may not be as effective in avian α2,3-linked NeuAc receptor binding and viral replication as WT avian virus but still show infection *in vivo*. Furthermore, reassortant viruses with an equine H7N7 HA and other genes from a chicken H5N2 IAV were shown to be lethal in chickens (11). This suggests a potential for transmission of equine-adapted viruses with NeuGc binding specificity back to avian species like chickens or ducks, for example, due to the close proximity of these species in farms. Our observations highlight the relevance of considering the fine receptor binding specificity when investigating the effect of species-specific adaptations in the RBS of HA and their potential in interspecies transmission events.

## Materials & Methods

### Cell culturing and preparation of cell lysates

The Biobank of the Friedrich-Loeffler-Institut (FLI), Greifswald Insel-Riems, Germany provided the following cell cultures for the *in vitro* characterization of the viruses: human embryonic kidney cells (HEK293T), Douglas Foster-1 cells (DF-1), Madin-Darby Canine Kidney type II cells (MDCKII), quail muscle 9 cells (QM-9), horse epidermal cells (E.Derm), horse lung cells (PLU-R), and duck embryo cells (SEF-R). In addition, 11d-old specific-pathogen-free embryonated chicken eggs SPF-ECE (Valo BioMedia, Germany) and chicken embryonic kidney cells (CEK) isolated from 18d-old SPF-ECE were used to perform replication kinetics (58).

MDCKII and PLU-R cells were cultured in minimal essential medium (MEM) with 10% fetal calf serum (FCS) containing Hank’s, Earls salts, NaHCO3, sodium pyruvate, and non-essential amino acids. For HEK293T, DF-1, QM-9, and SEF-R Iscove’s Modified Dulbecco’s medium (IMDM) with 10% FCS, Ham’s F12 nutrient mix, and glutamine was used. E.Derm and CEK cells were cultured in Eagle’s MEM and different concentrations of NaHCO3. HEK293S GnTI(-) cells were cultured in DMEM with 10% FCS. All cells were cultured at 37°C with 5% CO2.

MDCK WT (CCL-34) and MDCK-FUT (50) (a kind gift from Takahiro Hiono) cells were cultured in DMEM (Gibco) with 10% FCS (S7524, Sigma) and 1% penicillin and streptomycin (Sigma). MDCK-FUT cells were maintained with an additional 500 µg/ml G418 sulfate. MDCK-FUT cells overexpress the chicken fucosyltransferase genes *FUT3*, *FUT5*, and *FUT6* (50). Cells were cultured at 37°C with 5% CO2. Detaching of the cells was always done using 1X TrypLE Express Enzyme (12605010, Thermo Fisher Scientific), using 2 ml in a T75 flask, at a confluency of approximately 90%. Cell lysates were obtained using TrypLE Express Enzyme and RIPA lysis buffer as described previously (59).

### Viruses

The influenza viruses were obtained from different cooperation partners: A/chicken/Germany/R28/2003 (H7N7) (designated H7N7_avHA) was provided by Timm C. Harder, head of the reference laboratory for avian influenza virus, Friedrich-Loeffler-Institut (FLI), Greifswald Insel-Riems, Germany. AIV A/quail/California/D113023808/2012 (H4N2) was supplied by Beate Crossley from the California Animal Health and Food Safety Laboratory System, Department of Medicine and Epidemiology, University of California, Davis, United States. Stephan Pleschka and Ahmed Mostafa from Justus-Liebig-University, Gießen, Germany provided the human isolate A/Victoria/1975 (H3N2).

The avian sequence containing the five equine mutations S128T, I130V, A135E, T189A, and K193R (H3 numbering) was ordered from GenScript and inserted into the HA of A/chicken/Germany/R28/2003 in a pHWSccdB vector by restriction enzyme cloning. The IAV carrying these 5 mutations (designated H7N7_5eqHA) was generated in the backbone A/chicken/Germany/R28/2003 using reverse genetics. The virus was rescued in HEK293T cells, propagated in 9-to 11-d-old SPF eggs, and pooled for further use. Sequence analysis of different isolated viral pools revealed a stable establishment of the five mutations in the HA without a reversion to the WT sequence.

### α2,3-linked NeuAc receptor binding affinity assay

The binding of H7N7_avHA and H7N7_5eqHA to avian α2,3-NeuAc sialic acid receptor types was determined in a solid-phase binding assay using fetuin as a substrate as previously described (60, 61). The majority of sialic acids in fetuin are NeuAc and the low amount of NeuGc is neglectable for this assay (62). Briefly, 96 well-plates were coated with 200µl of 10µg/ml fetuin from fetal bovine serum (Merck, F3004) in 2x PBS overnight at 4°C. Fetuin-coated plates were washed with distilled water, dried at RT, and coated with 5 log^2^ HA units of H7N7_avHA, H7N7_5eqHA, A/quail/California/D113023808/2012 (H4N2) (positive control), or A/Victoria/1975 (H3N2) (negative control) in TBS at 4°C overnight. Viruses were tested in duplicates. Afterwards, plates were washed with PBS and then blocked for 1h RT using 0.1% de-sialylated BSA in 2x PBS. BSA was de-sialylated by incubating 5% BSA in 2x PBS + 0,02% penicillin-streptavidin with 1 unit of *Vibrio cholerae* neuraminidase for 24h at 60°C. The plate was washed with washing solution containing 2x PBS + 0.01% TWEEN80. Horseradish peroxidase (HRP) labeled fetuin was diluted 1:2 in 2x PBS with 0.02% TWEEN80 + 0.1% de-sialylated BSA and dilutions from 1:200 to 1:12,800 were added to the plate after washing and incubated for 1h at 4°C. After an additional washing step using washing solution, 100µl peroxidase substrate (Rockland; Lot# 24241) was added at RT. After 30 min the reaction was stopped using 50mM H2SO4 and the optical density was measured at 450 nm with an ELISA reader (Tecan).

### Plaque test and cell-to-cell spread

Plaque tests were performed using MDCKII for virus titration. Confluent MDCKII cells were incubated with diluted or undiluted samples for 1h at 37°C with 5% CO2. After infection, cells were washed twice using sterile PBS and overlaid with 1.8% bacto-agar in Dulbecco’s modified Eagle’s medium (DMEM) containing 5% bovine serum albumin (BSA). After incubation for 72 hours, plaque assays were fixed using a 10% formaldehyde solution with 0.1% crystal violet for 24h. After removal of the agar, the viral titers were calculated as PFU/ml or PFU/g, or the size of the plaques was measured under the microscope using Nikon NIS-Elements software.

### Replication kinetics

Different cell cultures were infected with H7N7_avHA and H7N7_5eqHA with a multiplicity of infection (MOI) of 0.001 for 1h. After washing with PBS, the cells were incubated at 37°C with 5% CO_2_. Cells and supernatants were harvested at indicated time points and stored at -80°C until the PFU/ml were assessed using plaque tests. The viral replication in SPF-ECE was tested in 11d-old eggs. Eggs were inoculated with 100 PFU/0.1 ml of each virus and incubated at 37°C with 56% humidity. Allantoic fluids were harvested at 8, 24, and 48 hpi and the PFU/ml was determined using plaque tests.

### Fusion assay to measure pH-dependent HA activation

The HA segments of H7N7_avHA and H7N7_5eqHA were cloned into the protein expression vector pCAGGS by restriction enzyme cloning (H7N7_avHA_pcAGGs and H7N7_5eqHA_pcAGGs respectively). A fusion assay was performed to assess the influence of the equine mutations on the pH activation of the HA as previously described (35). Briefly, confluent QM-9 cells were transfected in a 24-well plate with 500 ng of H7N7_avHA_pcAGGs and H7N7_5eqHA_pcAGGs and 100 ng GFP_pcAGGs per well using 1µl/µg DNA Lipofectamine 2000 (Thermofisher). PBS buffers were prepared with pH values ranging from 4.0 to 6.0 (0.2 steps). Transfected cells were incubated for 16h at 37°C with 5% CO_2_ and each well was washed with a different pH-adjusted PBS buffer for 10 min at RT after incubation. Cells were incubated for another 4h at 37°C with 5% CO_2_ and then fixed with 4% paraformaldehyde for 30 min at RT. The sizes of the fusion events were measured using a microscope and Nikon NIS-Elements software.

### Thermostability

The thermostability of H7N7_avHA and H7N7_5eqHA viruses was tested in a thermostability assay at 56°C. Allantoic fluid aliquots containing 10^5^ PFU/ml of viruses were prepared in tubes and incubated at 56°C. Samples were taken at 0, 1, 2, 4, and 6 hours post incubation. The infectivity of heat-exposed viruses was assessed in a plaque test on MDCKII cells. The PFU/ml are shown as mean values from two independent experimental rounds.

### Animal experiments

For the assessment of the pathogenicity index, nine chickens were infected intravenously (IV) and ten Pekin ducks were infected intramuscular (IM) with both recombinant viruses. Daily clinical scoring and the calculation of the pathogenicity index were performed according to the standard protocol of the world organization for animal health (WOAH). In addition, ten Pekin ducks and ten chickens were inoculated via the oculonasal route with 10^5^ PFU/ml. One day post-infection (dpi) five contact birds were added to each group to assess bird-to-bird transmission. In addition to the daily clinical scoring, oral and cloacal swab samples were taken on 2, 4, 7, and 10 dpi. The oral and cloacal swab samples were stored in MEM (H) + MEM (E) + NEA medium with BSA containing enrofloxacin, lincomycin, and gentamicin at -70°C until further use. The swab samples were titrated in plaque tests on MDCKII cells to assess the PFU/ml in the collected swabs. Organ samples were collected 4 dpi from three freshly dead or euthanized birds to assess viral distribution. Organ samples from the brain, heart, lung, liver, jejunum, spleen, and kidney were lysed in PBS using a tissue lyser and the PFU/g was assessed using plaque tests on MDCKII cells. The surviving birds were culled on day ten of the trial and blood was collected for serum samples. The serum was tested for influenza A NP antibody using a competitive ELISA kit (ID Screen Influenza A Antibody Competition Multispecies; IDvet).

### Expression and purification of HA for binding studies

HA encoding cDNAs of A/turkey/Italy/214845/02 H7N3 (63) (synthesized and codon-optimized by GenScript), A/duck/Australia/341/1983 H15N8 (a kind gift from Keita Matsuno), A/Vietnam/1203/2004 H5N1 (synthesized and codon-optimized by GenScript), A/chicken/Ibaraki/1/2005 H5N2 (43), and A/Taiwan/2/2013 H6N1 were cloned into the pCD5 expression vector as described previously (64). The pCD5 expression vector was adapted to clone the HA-encoding cDNAs in frame with DNA sequences coding for a secretion signal sequence, the Twin-Strep (WSHPQFEKGGGSGGGSWSHPQFEK); IBA, Germany), a GCN4 trimerization domain (RMKQIEDKIEEIESKQKKIENEIARIKK), and a superfolder GFP (65) or mOrange2 (66). Mutations in HAs were generated by site-directed mutagenesis (primers are available upon request). The HAs were purified from cell culture supernatants after expression in HEK293S GnTI(-) cells as described previously (64). In short, transfection was performed using the pCD5 expression vectors and polyethyleneimine I. The transfection mixtures were replaced at 6 h post-transfection by 293 SFM II expression medium (Gibco), supplemented with sodium bicarbonate (3.7 g/L), Primatone RL-UF (3.0 g/L, Kerry, NY, USA), glucose (2.0 g/L), glutaMAX (1%, Gibco), valproic acid (0.4 g/L) and DMSO (1.5%). At 5 to 6 days after transfection, tissue culture supernatants were collected and Strep-Tactin sepharose beads (IBA, Germany) were used to purify the HA proteins according to the manufacturer’s instructions.

### Glycan microarray binding of HA proteins

A selection of a glycan array that was presented elsewhere (67) was used and the full list of glycans is presented in Fig. S1. When fucosidase treatment was performed, cells were treated overnight at 37°C with 200 µg/ml fucosidase in fucosidase buffer (50 mM citrate buffer + 5 mM CaCl_2_, pH 6.0). Anti-sialyl-LewisX antibodies (mouse IgM, #551344, clone CSLEX1, BD Biosciences) at 50 µg/mL in 40 µL PBS-T were applied to the subarrays for 90 min. Subsequently, the arrays were incubated with a mixture of goat anti-mouse IgM HRP (10 µg/mL; #1021-05 Southern Biotech) and donkey anti-goat antibody labeled with AlexaFluor555 (5 µg/mL; A21432, Invitrogen) in 40 µL PBS-T for 1 h. HAs were pre-complexed with human anti-streptag and goat anti-human-AlexaFluor555 antibodies in a 4:2:1 molar ratio, respectively in 50 µL PBS-T on ice for 15 min. The mixtures were added to the subarrays for 90 min in a humidified chamber. Wash steps after each incubation (e.g. enzyme treatment, HA, or antibody incubation) involved six successive washes of the whole slides with either twice PBS-T, twice PBS, and twice deionized water. The arrays were dried by centrifugation and immediately scanned as described previously (27). Processing of the six replicates was performed by removing the highest and lowest replicates and subsequently calculating the mean value and standard deviation over the four remaining replicates.

### Fucosidase production

The protein sequence of fucosidase E1_10125 from *Ruminococcus gnavus* E1 (56) was obtained from the RCSB Protein Data Bank under accession number 6TR3. The nucleotide sequence was obtained from the closest hit in a protein-protein search in BLAST, of which the nucleotide sequence was corrected to obtain the exact same amino acid sequence. This open reading frame was ordered at GenScript and cloned into backbone pET23A in frame with a His-tag on the C-terminus of the open reading frame of the fucosidase. Cloning was performed in JM109 *Escherichia coli* (Promega). The plasmid is deposited at addgene under the name: pET23A-Fucosidase-E1_10125-Ruminococcus_gnavus-His (#207665). Expression of the fucosidase was performed in BL21 (DE3) *E. coli* (New England Biolabs). An overnight culture (90 ml per liter culture) was inoculated in 2xYT medium (Serva) supplemented with 50 µg/ml ampicillin (13398.02, Serva). Bacteria were grown at 37°C while shaking at 200 rpm until OD 0.8-1.0, after which fucosidase production was induced with 1mM isopropyl β-d-1-thiogalactopyranoside (R0309, Invitrogen). Afterward, the bacteria were grown for 21 h at room temperature while shaking at 200 rpm. Cell pellets were obtained by centrifugation in a swing-out centrifuge for 30 min at 4°C at 629 rcf. The pellets were resuspended in 50ml lysis buffer (100mM Tris-HCl, pH 8.0, 0.1%TritonX-100) per liter bacterial culture, which was supplemented with 1 gram per liter bacterial culture lysozyme (62971, Merck), 25 µl per liter bacterial culture DNAse (EN0521, Thermo Fisher Scientific), and 25 µl per liter bacterial culture DNAse buffer. The mixtures were incubated for 50 minutes at 37°C while shaking at 200 rpm. The cells were additionally lysed by sonication (Bandelin, Sonopuls, needle MS73) at 50% amplitude, three times for one minute at 10s intervals. The lysates were centrifuged for 1.5 h at 4°C at 629 rcf until the supernatant was clear. The supernatant was filtered through a 0.45 µm filter (431220, Corning) and incubated for 16h with Ni-NTA beads at 4°C while rotating. The beads were washed using a washing buffer (0.5M NaCl, 20mM Tris-HCl. pH 7.5) after which the fucosidase was eluted using the same buffer supplemented with 10-200mM imidazol. The elutions were concentrated and the buffer was exchanged to fucosidase buffer (50 mM citrate buffer + 5 mM CaCl_2_, pH 6.0) using a centrifugal concentrator with a molecular weight cutoff of 10 kDa (Vivaspin 6, VS0602, Sartorius). The presence of the fucosidase (62 kDa) was evaluated by running an SDS-PAGE gel (after denaturing for 15 min at 95°C with the addition of denaturing buffer (NP0009, Invitrogen)) with consequent staining using Coomassie blue dye.

### Protein histochemistry

Sections of formalin-fixed, paraffin-embedded chicken (*Gallus gallus domesticus*) were obtained from the Division of Pathology, Department of Biomolecular Health Sciences, Faculty of Veterinary Medicine of Utrecht University, the Netherlands. Sections of Pekin ducks were obtained from the animal experiment that is described above. Protein histochemistry was performed as previously described (68, 69). In short, tissue sections of 4 µm were deparaffinized and rehydrated, after which antigens were retrieved by heating the slides in 10 mM sodium citrate (pH 6.0) for 10 min. Endogenous peroxidase was inactivated using 1% hydrogen peroxide in MeOH for 30 min at RT. Slides were treated overnight at 37°C with 150 µg/ml fucosidase in the fucosidase buffer (50 mM citrate buffer + 5 mM CaCl2, pH 6.0) or buffer only. Subsequently, slides were washed with PBS with 0.1% Tween-20 (PBS-T). Tissues were blocked for 90 min at RT using 3% BSA (w/v) in PBS. Anti-sialyl-LewisX antibodies (mouse IgM, #551344, clone CSLEX1, BD Biosciences) were diluted 1:1000 in PBS and precomplexed with goat anti-mouse IgM-HRP (#1021-05, Southern Biotech) in a 1:100 dilution on ice for 20 min. The slides were stained for 90 min with pre-complexed HAs as previously described for the glycan microarray or the anti-sialyl-LewisX mixtures. For WT H5 HA, we used 1.5 µg/ml HA, for H5 R222K R227S HA, we used 3 µg/ml HA, for H7 HAs, we used 2 µg/ml HA, and for H15 HA, we used 1 µg/ml HA. After washing with PBS, binding was visualized using 3-amino-9-ethylcarbazole (AEC) (Sigma-Aldrich, Germany) and slides were counterstained using hematoxylin.

### Identification of *N*-glycans on cells by mass spectrometry

Cell lysates were obtained as described above. Preparation of mass spectrometry samples and measuring of these samples was performed as described previously (59). Briefly, glycans from cell lysates were released using PNGaseF treatment, labeled with procainamide 2-picoline borane, and purified in multiple steps. The samples were measured both using positive mode HILIC-IMS-QTOF analysis and MS/MS (using a Thermo Scientific Exploris 480 connected to a Thermo Scientific Ultimate 3000 UPLC system).

IM-MS data was calibrated to reference signals of m/z 121.050873 and 922.009798 using the IM-MS reprocessor utility of the Agilent Masshunter software. The mass-calibrated data was then demultiplexed using the PNNL preprocessor software using a 5-point moving average smoothing and interpolation of 3 drift bins. To find potential glycan hits in the processed data, the ‘find features’ (IMFE) option of the Agilent IM-MS browser was used with the following criteria: ‘Glycans’ isotope model, limited charge state to 5 and an ion intensity above 500. The found features were filtered by m/z range of 300 – 3200 and an abundance of over 500 (a.u.) where abundance for a feature was defined as ‘max ion volume’ (the peak area of the most abundant ion for that feature). After exporting the list of filtered features, glycans with a mass below 1129 Da (the mass of an *N*-glycan core) were removed. The ExPASy GlycoMod tool (70) was used to search for glycan structures (monoisotopic mass values, 5 ppm mass tolerance, neutral, derivatized N-linked oligosaccharides, procainamide (mass 235.168462) as reducing terminal derivative, looking for underivatized monosaccharide residues (Hexose, HexNAc, Deoxyhexose, and NeuAc)). For features with multiple potential monosaccharide combinations, the most realistic glycan in the biological context was chosen. The abundance of glycan features with the same mass, composition, and a maximum difference of 0.1 min in the retention time were combined as one isomer. A full glycan composition feature list for MDCK-FUT cells is presented in Table S2. Analysis of the number of fucoses per sialylated glycan was performed on the complex and hybrid *N*-glycans with at least one sialic acid.

For MS/MS data, Proteowizard MSconvert (version 3.0.21328-404bcf1) was used to convert Thermo raw files to MGF format using MGF as output format, 64-bit binary encoding precision and with the following options selected: write idex, zlib compression and TPP compatibility. No filters were used when converting raw files to MGF format. To search MGF files for spectra containing glycan oxonium ions an internally developed tool named Peaksuite (v1.10.1) was used with an ion delta of 20 ppm, noise filter of 0% and using a list of oxonium *m/z* values as mass targets (Table SX from (59)). Scans without any detected peaks were removed. Python 3.2.2 was used for data curation based on precursor *m/z* (10 ppm), retention time (17-24 min) and intensities of oxonium ions that originated from the glycan core (*m/z* 441.2707, 587.3286, 644,3501, 790.4080, 806.4029, and 952.4608). The sum intensity threshold of the core oxonium ions was set to 1e4. Python 3.2.2 was also used for calculating the relative intensities of oxonium ions corresponding to sLe^X^ (*m/z* 803.2928) normalized versus the sum intensities of the core oxonium ions.

### Flow cytometry studies

Detaching of the cells (MDCK WT and MDCK-FUT) was performed with 1X TrypLE Express Enzyme (12605010, Thermo Fisher Scientific), using 2 ml in a T75 flask, at a confluency of approximately 90%. Cell pellets were obtained by centrifugation for 5 min at 250 rcf, after which the cells were resuspended in PBS supplemented with 1% FCS (S7524, Sigma) and 2mM EDTA and kept at 4°C until the plate was measured in the flow cytometer. In a round-bottom 96-well plate (353910, Falcon), 50,000 cells per well were used. Per well, 1 µg of precomplexed HA or precomplexed anti-sialyl-LewisX antibody (CD15S, clone CSLEX1, #551344, BD Biosciences) was added to PBS supplemented with 1% FCS and 2mM EDTA to achieve a final concentration of 20 µg/ml. Precomplexation of HAs was performed with 1.3 µg monoclonal antibody detecting the Twin-Strep-tag and 0.325 µg goat anti-human Alexa Fluor 488 (A11013, Invitrogen) and incubated on ice for 20 min. Precomplexation of the anti-sialyl-LewisX antibody was performed with a 1:50 dilution of goat anti-mouse IgM-HRP (#1021-05, Southern Biotech) and 0.65 µg donkey anti-goat Alexa Fluor 555 (A21432, Invitrogen) and incubated on ice for 20 min. Furthermore, eBioscience Fixable Viability Dye eFluor 780 (65-0865, Thermo Fisher Scientific) was diluted 1:2000 in the same mixture. Cells were incubated with the hemagglutinin/antibody mix for 30 minutes at 4°C in the dark.

Cells were washed once with PBS supplemented with 1% FCS and 2 mM EDTA, after which the cells were fixed with 100 µl of 1% paraformaldehyde in PBS for 10 minutes. Afterward, cells were washed twice using PBS supplemented with 1% FCS and 2 mM EDTA, after which they were resuspended in 100 µl of the same buffer. Flow cytometry was performed using the BD FACSCanto II (BD Biosciences) using appropriate laser voltages. Alexa Fluor 488 signal (HAs) was measured using the FITC filter and Alexa Fluor 555 signal (anti-sialyl-LewisX signal) was measured using the PE filter. Data were analyzed using FlowLogic (Inivai Technologies) and gated for cells, single cells, and alive cells as described previously (59). The mean value and standard deviation were calculated over triplicate measurements.

### Sequence analysis

The prevalence of the five equine amino acid residues on positions 128, 130, 135, 189, and 193 (numbering according to H3 HA) was analyzed in all equine and avian H7 sequences with a minimum length of 1,600 bp. The protein sequences were downloaded from GISAID (date of download: 31.08.2023) and aligned using MAFFT package in Geneious software. Only the receptor binding site of the different sequences was compared. In total 3,402 avian H7 and 24 equine H7 sequences were analyzed.

### Data analysis and statistical analysis

The data in this article were (statistically) analyzed and visualized using GraphPad Prism 9.2.0.

## Data deposition

The released *N*-glycan raw data (for the glycomic analyses of the cell lines) have been deposited to the GlycoPOST repository (Watanabe, Y., Aoki-Kinoshita, K.F., et al. 2021) under the identifier GPST000345.

## Acknowledgments

We would like to thank Dajana Helke, Nadine Bock, Luise Hohensee, and Bastiaan Vroege for laboratory technical assistance. Furthermore, we thank the animal caretakers and the department of experimental animal facilities and biorisk management at the FLI-Riems for their support in the animal experiments. Timm C. Harder, Ahmed Mostafa, Stephan Pleschka, and Beate Crossley are thanked for providing the viruses, and Stefan Finke for providing the pCAGGS plasmid used in this study. We thank the authors who submitted nucleotide sequences in the GISAID that were used for analysis in this study.

## Ethics statement

All performed animal experiments were approved and monitored by the commissioner for animal welfare at the Friedrich-Loeffler-Institut (FLI), representing the Institutional Animal Care and Use Committee (IACUC). The animal trial was performed under biosafety level 3 (BSL3) conditions in the animal facilities of FLI according to the German Regulations for Animal Welfare. The experiments were approved by the authorized ethics committee of the State Office of Agriculture, Food Safety, and Fishery in Mecklenburg-Western Pomerania (LALLF M-V) under registration number 7221.3–1.1-051-12. The specific-pathogen-free (SPF) embryonated chicken eggs (ECE) used for this publication were purchased from Valo BioMedia (Germany). The storage and handling of SPF-ECE were performed according to the guidelines of the World Organization for Animal Health (WOAH).

## Funding

This research was made possible by funding from ICRAD, an ERA-NET co-funded under the European Union’s Horizon 2020 research and innovation programme (https://ec.europa.eu/programmes/horizon2020/en), under Grant Agreement n°862605 (Flu-Switch) to R.P. de Vries and E.M. Abdelwhab. This work was also supported by grants from the Deutsche Forschungsgemeinschaft (DFG, AB 567, to E.M. Abdelwhab), European Commission (ERC Starting Grant 802780 to R.P. de Vries), and the Mizutani Foundation for Glycoscience (Research Grant 2023 to R.P. de Vries).

## Supplementary information

**Table S1.**
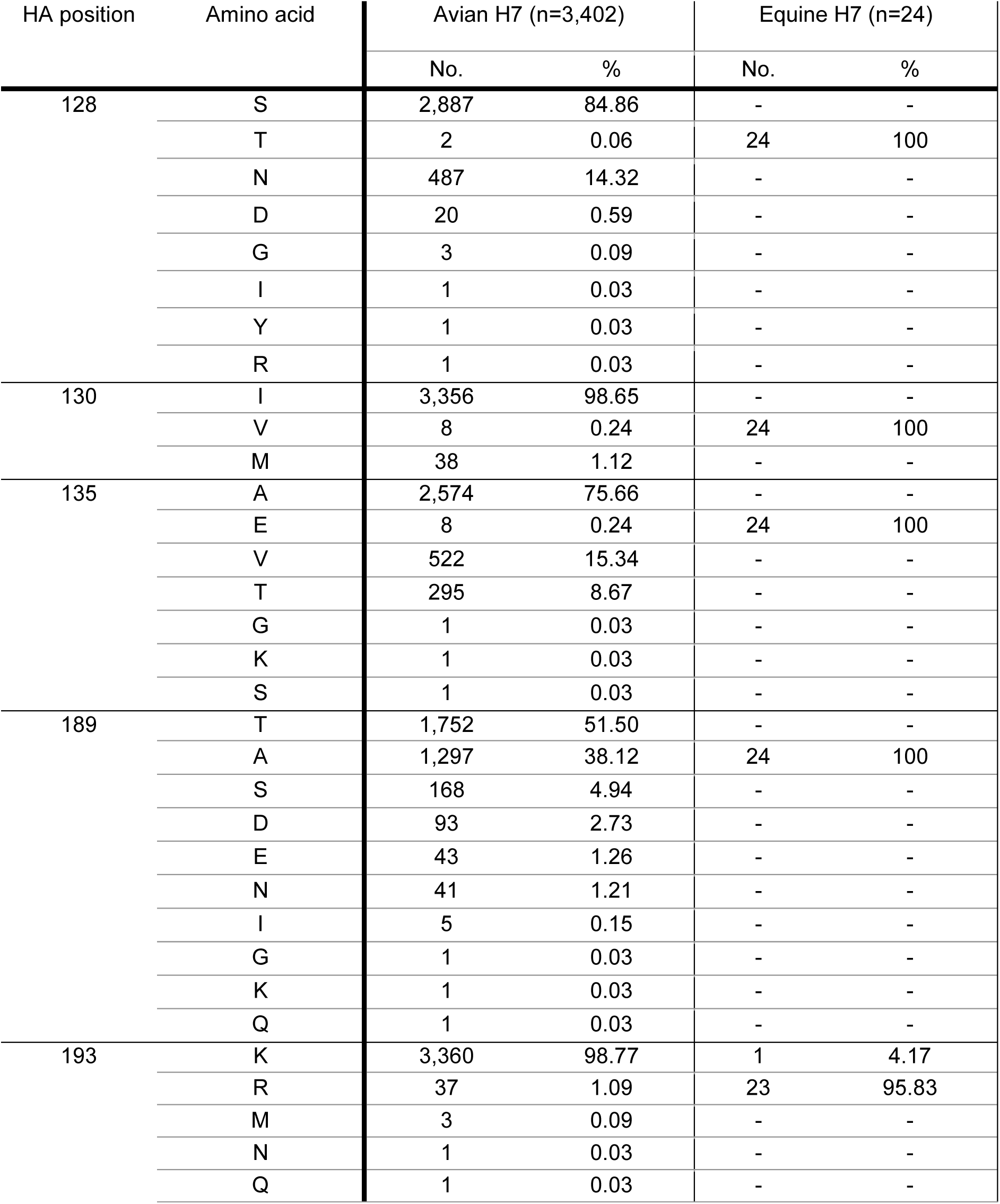
Prevalence of different amino acid residues in avian and equine H7 HA sequences. Sequences with a size of at least 1,600bp were downloaded from GISAID on 31.08.2023 and analyzed using Geneious software.

**Table S2. Relative abundance of *N*-glycans of MDCK FUT cells measured using HILIC-IMS-QTOF positive mode mass spectrometry.** The table is presented in an additional excel file. The data from MDCK WT cells is published in Table SII of (59).

**Figure S1.**
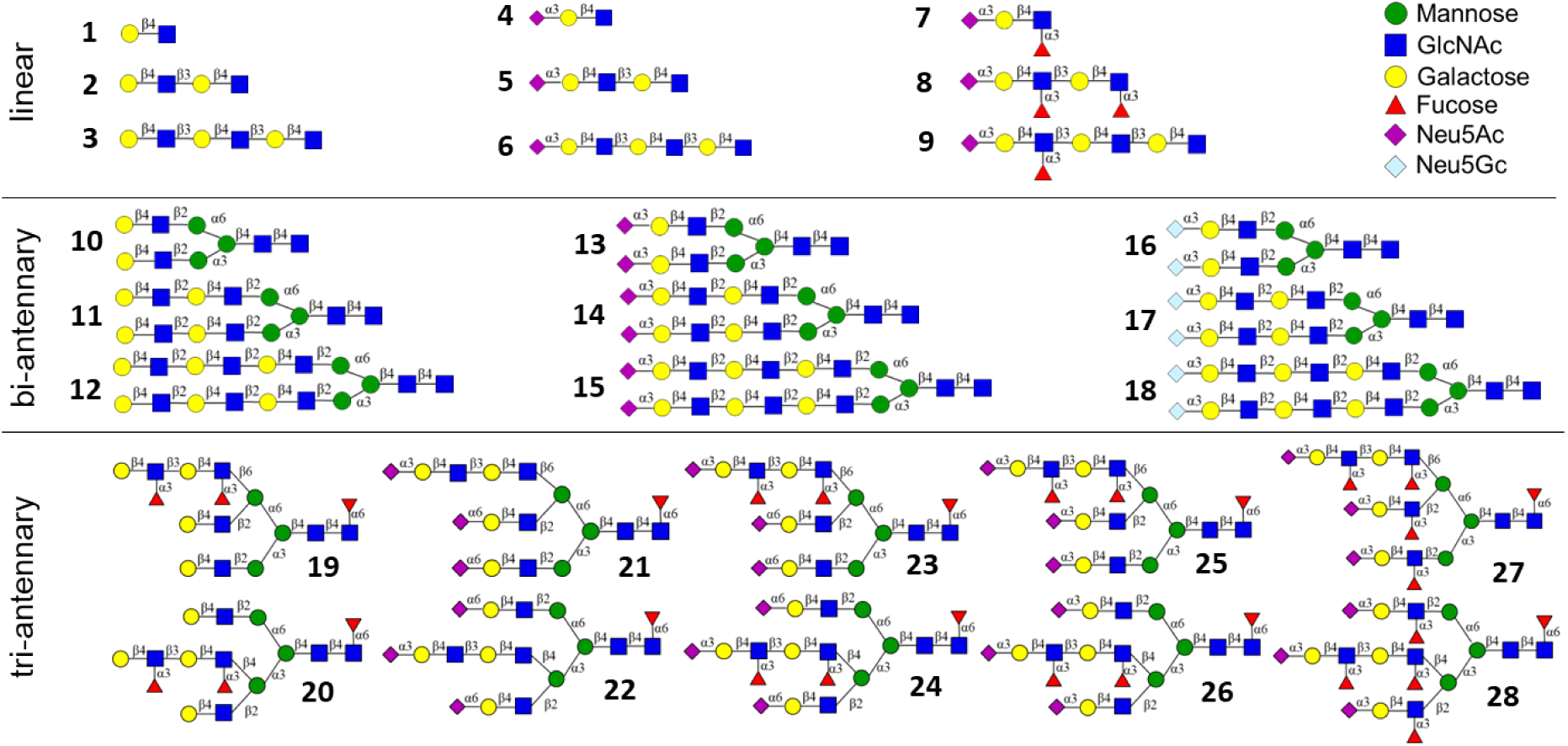
Glycans presented on the glycan microarray

